# Iron activates microglia and directly stimulates indoleamine-2,3-dioxygenase activity in the N171-82Q mouse model of Huntington’s disease

**DOI:** 10.1101/550905

**Authors:** David W. Donley, Marley Realing, Jason P. Gigley, Jonathan H. Fox

## Abstract

Huntington’s disease (HD) is a neurodegenerative disorder caused by a dominant CAG-repeat expansion in the huntingtin gene. Morphologic activation of microglia is a key marker of neuroinflammation that is present before clinical onset in HD patients. The kynurenine pathway of tryptophan degradation is restricted in part to microglia and is activated in HD, where it contributes to disease progression. Indoleamine-2,3-dioxygenase (IDO) is a microglial enzyme that catalyzes the first step in this pathway. HD brain microglial cells also accumulate iron; however, the role of iron in promoting microglial activation and the kynurenine pathway is unclear. Based on analyses of morphological characteristics of microglia, we showed that HD mice demonstrate an activated microglial morphology compared with controls. Neonatal iron supplementation resulted in additional microglial morphology changes compared with HD controls. Increased microglial activation in iron-supplemented HD mice was indicated by increased soma volume and decreased process length. In our assessment of whether iron can affect the kynurenine pathway, iron directly enhanced the activity of human recombinant IDO1 with an EC_50_ of 1.24 nM. We also detected elevated microglial cytoplasmic labile iron in N171-82Q HD mice, an increase that is consistent with the cellular location of IDO. We further demonstrated that neonatal iron supplementation, a model for studying the role of iron in neurodegeneration, activates IDO directly in the mouse brain and promotes neurodegeneration in HD mice. Kynurenine pathway metabolites were also modified in HD and by iron supplementation in wild-type mice. These findings indicate that iron dysregulation contributes to the activation of microglia and the kynurenine pathway in a mouse model of HD.

## Introduction

Huntington’s disease (HD) is an autosomal-dominant neurodegenerative disorder caused by a CAG-repeat expansion in exon 1 of the huntingtin gene (*HTT*) that results in expression of mutant huntingtin protein (mHTT) (HDCRG, 1993). Mutant HTT is expressed throughout the central nervous system; however, the striatum and cortex are the main brain areas that degenerate in HD. Mutant HTT accumulates and misfolds in neurons and also microglial cells (H.P. et al., 2017). Multiple mechanisms are implicated in the neurodegeneration that occurs in HD. Expression of mHTT results in altered transcription, axonal transport, mitochondrial function, vesicular trafficking, inflammation, and oxidative stress (La Spada et al., 2011). Exactly how these pathways are related is not fully understood, but they result in progressive neuronal dysfunction and loss, eventually leading to disease onset and clinical progression. Symptoms of HD encompass a hyperkinetic motor disorder, cognitive decline, and varied psychiatric problems (Vonsattel and DiFiglia, 1998). Currently, there are no interventions that delay the onset or progression of HD.

Numerous studies have shown that iron accumulates in the brains of HD patients. Magnetic resonance imaging–based approaches have provided evidence that iron accumulates progressively, starting before clinical onset and occurs in a manner dependent on CAG expansion size (Bartzokis et al., 1999, Vymazal et al., 2007, Jurgens et al., 2010). Significant iron accumulation in HD patients occurs in the striatum and cerebral cortex, regions that show early degeneration, suggesting that iron accumulation may have a role in neurodegeneration (Rosas et al., 2012, Bartzokis et al., 1999). Post-mortem studies also demonstrate increases in regional iron levels in brains from individuals with advanced HD (Rosas et al., 2012, van Bergen et al., 2016, Dexter et al., 1991). However, the approaches used in these studies could not determine the brain cell types involved and do not reliably differentiate between the accumulation of iron in a safe form, such as ferritin-bound, and in a potentially toxic labile form. Thus it is unclear how accumulated iron is contributing to HD pathogenesis.

Huntingtin protein is involved in iron homeostasis (Dietrich et al., 2017, Lumsden et al., 2007), but the mechanisms through which mHTT results in brain iron accumulation are incompletely understood. Microglia have an important role in brain iron metabolism (Andersen et al., 2014). Being phagocytic, they help to clear dying cells, which can result in their uptake of iron (Yoshida et al., 1995, Rathnasamy et al., 2011). In addition, as the primary brain immune cell, microglial cells also accumulate iron in response to inflammation (Lopes et al., 2008, McCarthy et al., 2018, Castelnau et al., 1998). Although the mechanisms involved are not fully understood, microglial iron accumulation is present in human HD brains and is associated with an activated microglial morphology (Simmons Danielle et al., 2007). Further, HD mouse models demonstrate increased ferritin-bound iron, and microglia have increased ferritin expression and detectable iron(III) stores (Simmons Danielle et al., 2007, Chen et al., 2013). Iron within the ferritin shell is considered a safe form, as it is not able to interact with oxygen. Therefore, these findings suggest that microglia in HD demonstrate a protective response to brain iron accumulation and degeneration. However, it is unclear if this response is complete or if there is additional accumulation of labile iron in microglial cells that promotes inflammatory or degenerative processes. Microglial iron promotes oxygen radical generation that potentiates inflammation and neurotoxicity in models of Parkinson’s disease (Zhang et al., 2014, Mehlhase et al., 2006); however, it is unknown if iron dysregulation promotes inflammation in mouse models of HD.

The kynurenine pathway (KP) of tryptophan metabolism is involved in the progression of HD and occurs, in part, in microglial cells (Beal et al., 1990, Guidetti et al., 2000). This pathway is activated by inflammatory cytokines, including IL-6, that are upregulated early in HD (Björkqvist et al., 2008). The first step in the KP is the oxidation of tryptophan to kynurenine by the enzyme indoleamine-2,3-dioxygenase (IDO). In the brain, IDO is upregulated by microglial activation (Yadav et al., 2007, Corona et al., 2013). KP activity in HD is thought to contribute to disease progression via increased flux through a pathway branch that generates neurotoxic intermediates including 3-hydroxykynurenine (3-HK) and decreased flux through a branch that generates protective kynurenic acid (Campesan et al., 2011b, Beal et al., 1990, Guidetti et al., 2000, Guidetti et al., 2004, Zwilling et al., 2011). Iron modifies KP activity by activating 3-hydroxyanthanilic acid dioxygenase (3-HAO) and promoting synthesis of neurotoxic quinolinic acid (Stachowski and Schwarcz, 2012). Despite this, it is unknown if iron has a key regulatory role in the KP and if it modulates gatekeeping steps such as IDO-mediated oxidation of tryptophan.

Even though HD is caused only by the CAG expansion of the huntingtin gene, an estimated 60% of the non-CAG variance in age of onset is explained by as-yet-unknown environmental factors (Wexler, 2004). We have previously used neonatal iron supplementation (NIS) to assess the possible role of high dietary iron intake as an environmental modifier of HD in mice. This model is relevant, as in most populations human infants are supplemented with iron to prevent deficiency. Recommended daily allowances for iron have been developed to prevent iron deficiency, and upper limits were established to avoid short-term adverse effects (Micronutrients, 2001); however, long-term consequences of iron supplementation are not understood (Agrawal et al., 2017). We have previously shown that NIS in R6/2 and YAC128 mouse models of HD exacerbates disease progression (Berggren et al., 2015, Berggren et al., 2016). We have additionally shown that NIS promotes markers of mitochondrial dysfunction in R6/2 mice (Agrawal et al., 2018), although the cell-specific mechanisms by which NIS promotes HD in mice are unknown.

HD patients develop presymptomatic microglial activation and elevated blood cytokine levels (Björkqvist et al., 2008, Tai et al., 2007, Sapp et al., 2001). Activated microglia display morphological changes that correlate with increased production of proinflammatory cytokines such as IL-1β and increased phagocytic activity (Fernández-Arjona et al., 2017, Davis et al., 2017, Sapp et al., 2001). Cytokines including IL-10, TNFα, IL-8, and IL-6 are elevated in blood presymptomatically and correlate with disease severity (Björkqvist et al., 2008). In HD mouse models, the detrimental role of inflammation is supported by reports of adverse effects of proinflammatory responses to lipopolysaccharide and the protozoal pathogen *Toxoplasma gondii* (Björkqvist et al., 2008, Donley et al., 2016). In addition, Laquinimod, a drug that reduces inflammation, is protective in HD mice (Garcia-Miralles et al., 2016). Despite evidence for roles of iron dysregulation and microglial-mediated inflammation in HD, it is unclear if these pathways interact. Here we examined the role of iron in promoting microglial and KP activation in the N171-82Q mouse model of HD. We used the NIS paradigm to examine effects on HD outcomes, as well as morphologic and KP measures of microglial activation. We report that iron directly activates IDO at physiologically relevant concentrations, differentially activates microglial cells in HD versus wild-type (WT) mice following neonatal supplementation, and accumulates in labile form in HD microglia. Our findings support the involvement of iron in microglial activation, particularly in the context of early-life supplementation. This study has relevance for understanding how dysregulation of brain iron homeostasis promotes HD and also how increased dietary iron intake potentiates HD in mouse models.

## Results

### Elevated indoleamine-2,3-dioxygenase activity in HD mouse brain is decreased by iron chelation

We measured brain IDO activity in the striatum and cortex of WT and HD mice with and without NIS. IDO activity was significantly increased in striata (F_(1,42)_ = 7.77, p = 0.0075) and cortices (F_(1,42)_ = 13.57, p < 0.001) of HD mice regardless of NIS (**Fig. 1A,B**). In HD but not WT cortices, NIS significantly increased IDO activity (p = 0.016). IDO activity in HD mice that received NIS was significantly decreased by *ex vivo* deferoxamine addition in the striata (p = 0.0067) and in the cortices (p = 0.0028) (**Fig. 1A,B**). IDO has two isoforms, IDO1 and IDO2. IDO1 is highly expressed in brain microglia as well as in peripheral tissues and is a regulator of KP activity in the context of inflammation (Schwarcz et al., 2012, Lee et al., 2014, Ball et al., 2007). The second isoform, IDO2, is more narrowly expressed, primarily in peripheral tissues, and has much lower activity as compared with IDO1 (Metz et al., 2007, Ball et al., 2007). Therefore, we measured *Ido1* and *Ido2* transcript levels relative to beta-actin in WT and HD brains. *Ido1* was not significantly increased in the striatum (F_(1,20)_ = 1.36, p = 0.2581) but was increased in the cortex of HD mice (F_(1,20)_ = 5.86, p = 0.0251) relative to WT mice regardless of NIS (**Fig. 1C,D**). We found in both the striatum (F_(1,20)_ = 4.42, p = 0.0484) and cortex (F_(1,20)_ = 4.82, p = 0.0402) that NIS alone significantly increased *Ido2* expression irrespective of mouse genotype (**Fig. 1E,F**). Together, these data demonstrate that *Ido* expression does not account for all the changes in enzyme activity and that iron may have a direct role in activating enzyme activity.

**Figure 1:**
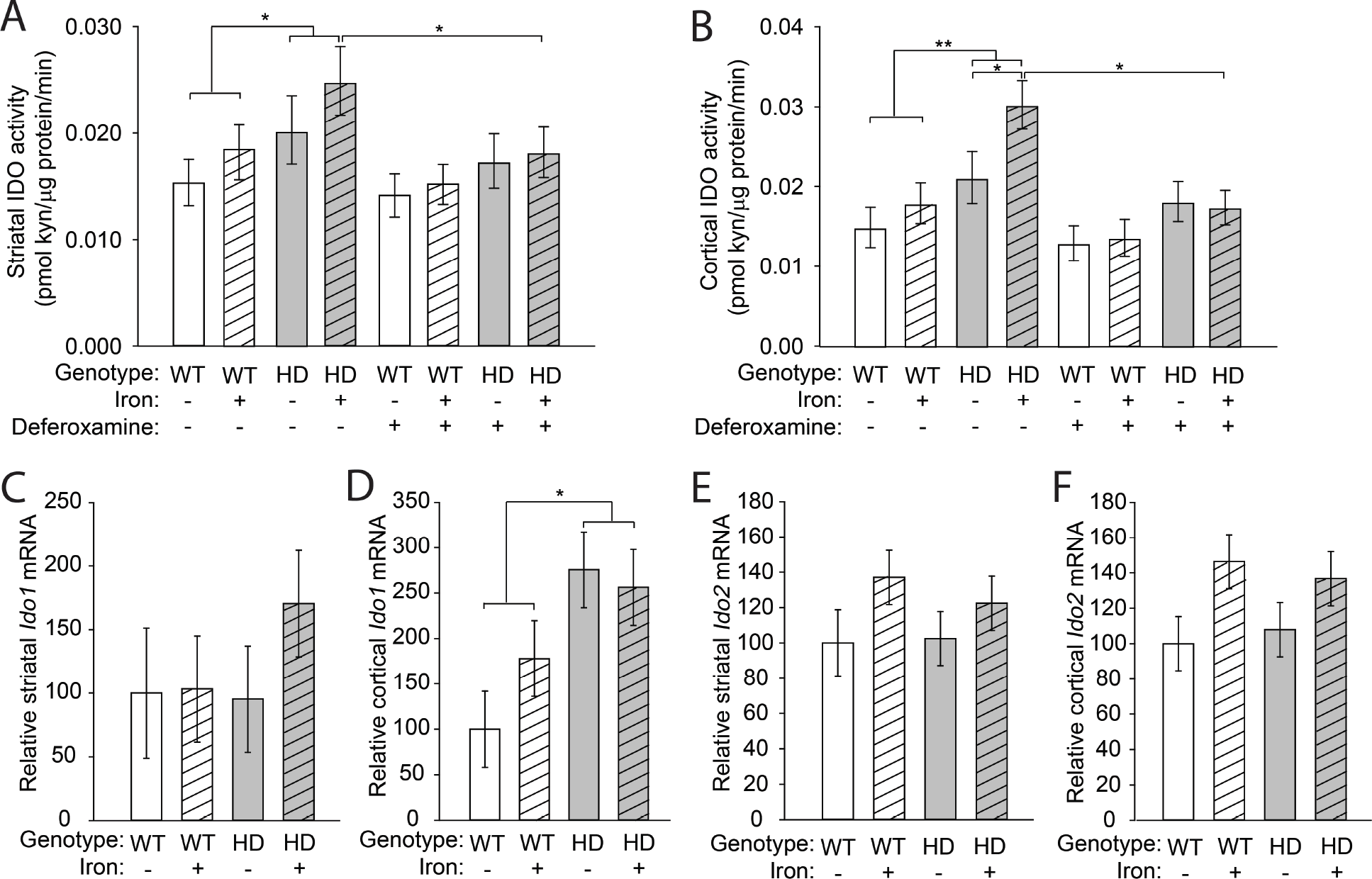
Elevated IDO activity in HD mouse brains is decreased by *ex vivo* iron chelation. Mouse pups were supplemented with carbonyl iron from postnatal days 10–17 and then were sacrificed at 14 weeks of age. **A.** Striatal IDO activity is increased in N171-82Q HD mice and significantly decreased by *ex vivo* iron chelation. **B.** Cortical IDO activity is increased in iron-supplemented HD mice compared to non-supplemented HD mice and activity is decreased by 50 μM deferoxamine. **C**. There is no difference in relative transcript levels of IDO1 in striatum. **D.** IDO1 mRNA is increased in cortex of HD mice at 14-weeks of age. **E,F**. Iron increases levels of IDO2 in striatum (**E**) and cortex (**F**) but there was no significant effect of HD on IDO2 mRNA. **A,B**. Bars represent means ± 95% CI. n=9 wild-type and 11 HD mice in each group. *p<0.05, **p<0.01. **C,F**. Bars represent means ± SE, n=6; *p<0.05.

### Iron directly activates IDO

To test whether iron stimulates IDO enzymatic activity, we incubated mouse brain extracts with iron(II). Iron increased brain IDO activity (F_(3,16)_ = 14.00, p < 0.001) in a dose-dependent manner (**Fig. 2A**). To determine if iron directly activates IDO, we measured the activity of purified human IDO1 enzyme in context of iron. We again measured a dose-dependent increase in IDO activity (F_(5,10)_ = 6.42, p = 0.0064) with an EC_50_ = 1.24 nM for iron activation of IDO1 (**Fig. 2B**). We then studied the effect of increased iron on cultured mouse microglial cells. After 2 hours of iron incubation in serum-free medium, we measured significantly increased IDO activity (F_(6,14)_ = 6.04, p = 0.0027) (**Fig. 2C**). To study whether this effect was a post-translational mechanism, we supplemented iron into microglial cultures in the presence of the protein translation inhibitor cycloheximide. We performed this experiment in serum-containing medium to reduce toxicity due to protein translation inhibition. Iron addition significantly increased IDO activity (F_(3,19)_ = 14.11, p < 0.001); however, we found no difference in IDO activity with cycloheximide treatment, supporting a direct, post-translational activation of IDO by iron (**Fig. 2D**).

**Figure 2:**
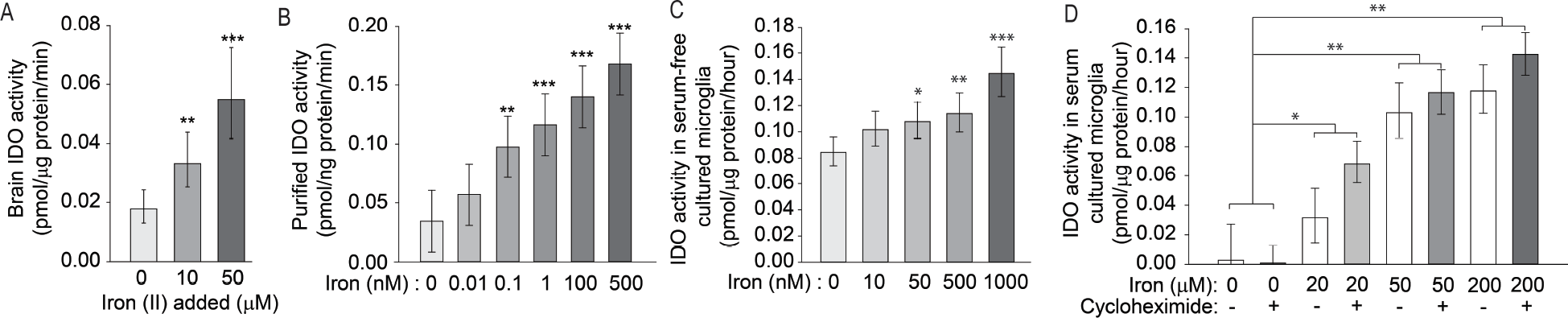
Nanomolar iron activates IDO. The effect of iron on IDO activity was determined in brain extracts, purified protein, and cultured EOC20 mouse microglial cells. **A**. Ferrous ammonium sulfate increased IDO activity in protein extracts from 14-week-old WT mouse brains in a dose-dependent manner. n=5. **B**. Ferrous iron activated purified IDO. n=3. **C**. Iron(II) added to cultured microglia in serum-free medium for 2 hours increased IDO activity. n=3. **D**. Iron(II) was added to the medium of cultured microglia for 2 hours with and without addition of cycloheximide, and IDO activity was determined. IDO activity was significantly increased by iron addition but no difference was seen with addition of cycloheximide. n=6. **A,D**. Data are shown as the mean ± 95% CI. **B,C**. Data are shown as the mean ± SE. *p<0.05, **p<0.01, ***p<0.001. **A-C.** Asterisks represent comparisons to samples without iron addition.

### Neonatal iron supplementation exacerbates neurodegeneration in N171-82Q HD mice

NIS in R6/2 and YAC128 HD mouse models exacerbates neurodegeneration (Berggren et al., 2015, Berggren et al., 2016). Consistent with this, both N171-82Q HD mice (F_(1,61)_ = 77.70, p < 0.001) and NIS (F_(1,61)_ = 4.28, p = 0.0428) significantly decreased brain weights (**Fig. 3A**). There was a significant interaction (F_(1,61)_ = 4.05, p = 0.0486) between HD and NIS (**Fig. 3A**); iron-supplemented HD mice had significantly decreased brain weights compared with control HD mice (p = 0.0072) and iron-supplemented WT mice (p < 0.001). In addition, striatal volumes were significantly decreased in HD mice (F_(1,30)_ = 10.32, p = 0.0031), and there was a trend toward a genotype-treatment interaction (F_(1,30)_ = 3.53, p = 0.0702) (**Fig. 3B**). No difference in striatal volume was found comparing control HD mice with control WT mice (p = 0.3559). However, iron-supplemented HD mice had significantly decreased striatal volume compared with control HD mice (p = 0.0206) and with iron-supplemented WT mice (p=0.0011). There was no difference among groups with respect to striatal neuron cell body volume, suggesting neuronal loss is occuring (**Fig. 3C**). We also characterized mouse behavioral phenotypes using rotarod endurance testing and body weight measured over time. Consistent with previous reports in this model (Lu et al., 2014, Masuda et al., 2008), HD mice had poorer motor performance on the rotarod and decreased body weights by 10 weeks of age. There was no effect of NIS on these outcomes (data not shown).

**Figure 3:**
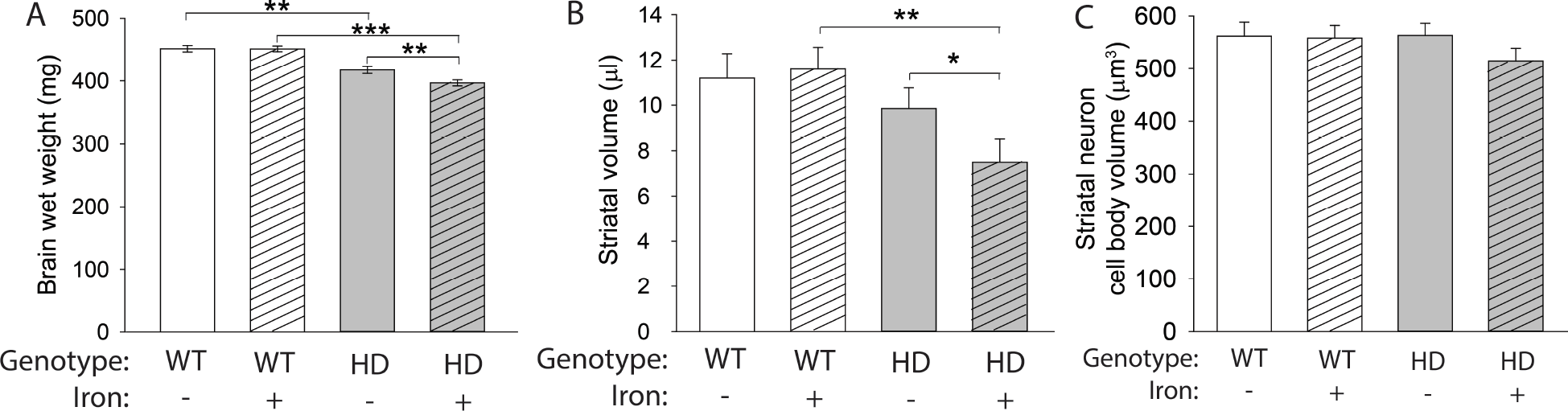
Neonatal iron supplementation exacerbates HD in the N171-82Q mouse model. Mouse pups were administered neonatal iron from postnatal days 10–17 then were sacrificed at 14 weeks of age. **A**. Total brain wet weight decreased in HD mice at 14 weeks of age. n=15 WT control, 19 WT NIS, 13 HD control, 17 HD NIS. **B**. Striatal volume measured using the Cavalieri method was decreased in iron-supplemented HD mice compared with both control HD mice and iron-supplemented WT mice. **C**. Cell body volume of striatal neurons was not significantly different among groups. Data are shown as the mean ± SE. **p<0.01, ***p<0.001. **B,C.** n=10 WT control, 11 WT NIS, 12 HD control, 11 HD NIS.

### Neonatal iron supplementation induces an activated microglial morphology in HD mice

Activated microglia demonstrate altered morphology and are present in HD (Kraft et al., 2012, Simmons Danielle et al., 2007). We therefore quantified microglial morphology as an indicator of activation status. Microglia were characterized as ramified, primed, amoeboid, or reactive (Kraft et al., 2012, Torres-Platas et al., 2014) (**Fig. 4A**). To address whether iron supplementation and/or mouse genotype is associated with microglial activation as assessed by their morphology, we performed logistic regression. This test allowed us to statistically test differences in the blinded, categorical assignment of microglial activation. Ramified cells are considered to be non-activated and perform a different function than activated microglia whereas the primed morphology is an intermediate functional and morphological phenotype between the non-activated ramified cells and the highly activated amoeboid and reactive microglia (Kierdorf and Prinz, 2013, Lawson et al., 1990). Therefore, we first compared the proportion of ramified microglia to the combined proportion of primed, amoeboid, and reactive cells. We found that iron significantly increased the probability of an activated microglial morphology. The proportion of ramified and primed microglia was then compared to highly activated amoeboid and reactive microglia to determine the probability of a microglial cell exhibiting a fully activated morphology. We again found that iron significantly increased microglial activation and that HD mice had a higher proportion of highly activated microglia (amoeboid or reactive) when supplemented with iron (p-values in **Fig. 4B**). The percentages of the four morphologies relative to total microglial cells are illustrated in **Fig. 4B** for comparison purposes. To extend these findings, we also quantified morphological characteristics using a quantitative neurolucida-based approach. NIS increased cell body volumes (F_(1,124)_ = 8.74, p = 0.0037), but there was no difference between iron-supplemented WT and HD microglia (**Fig. 4C**). We also found a significant NIS-HD interaction with respect to the number of microglial processes (F_(3,40)_ = 4.8, p = 0.006); these were significantly increased in iron-supplemented HD mice relative to control HD mice (p = 0.0024) (**Fig. 4D**). In contrast, HD mice had decreased process lengths (F_(1,124)_ = 6.09, p = 0.0149), which were unaffected by iron supplementation (**Fig. 4E**).

**Figure 4:**
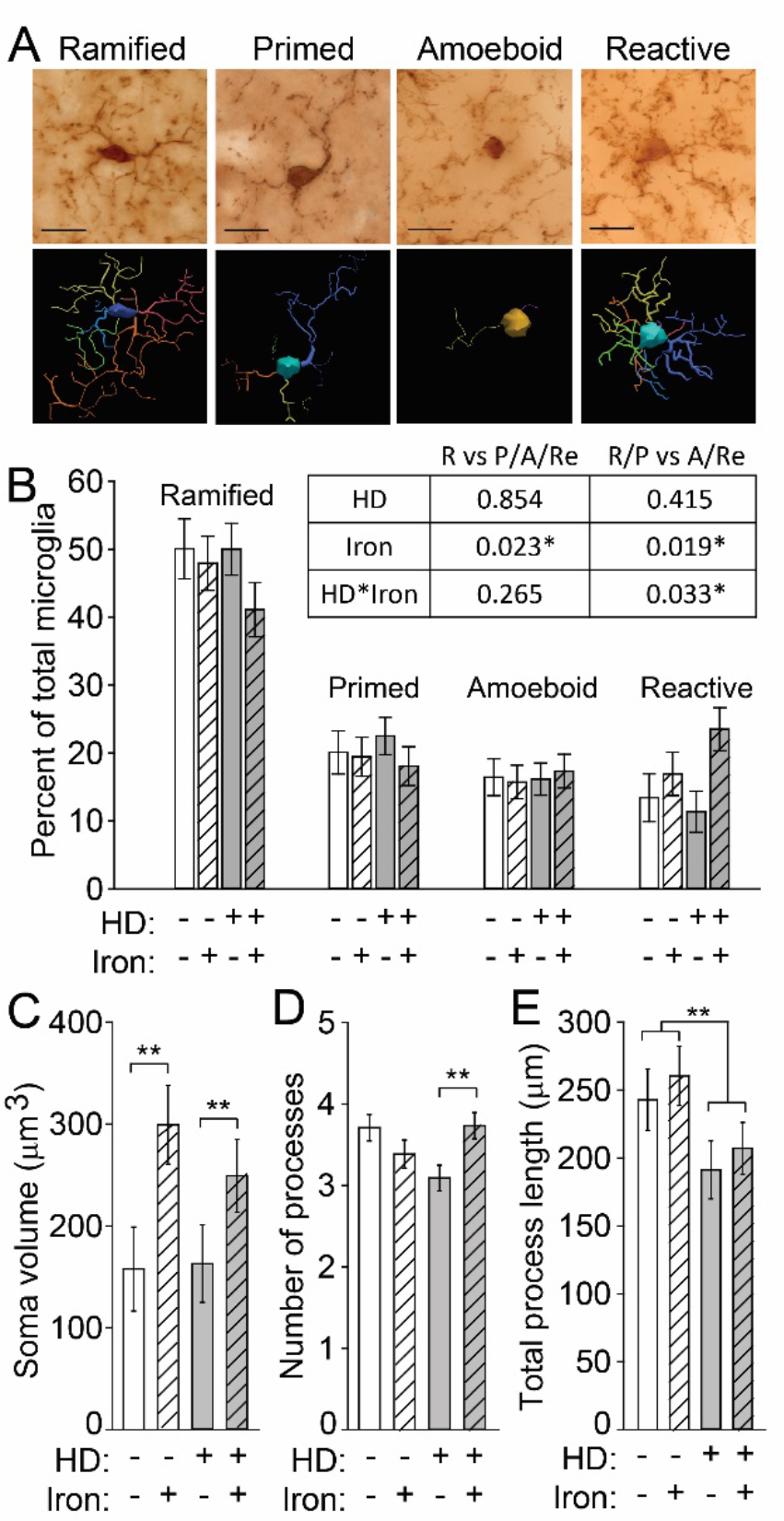
Neonatal iron supplementation and HD differentially alter microglial morphology. Microglia in brain sections from 14-week-old N171-82Q mice were labeled with anti-Iba1, and their morphology was evaluated. **A**. Representative micrographs of microglial cells labeled with anti-Iba1 were identified and characterized as having one of four morphologies, amoeboid, primed, ramified, and reactive. Neurolucida traces are shown below each image. Scale bars = 20 μm. **B.** Iron increased the frequency of reactive microglia in iron-supplemented HD mice. The table shows p-values from logistic regression analysis of microglial morphologies. Iron significantly increased the probability of activated microglial morphologies. R=ramified; P=primed; A=amoeboid; Re=reactive. Each morphology is presented as a percent of total microglial cells within each mouse. The percentage of cells with each morphology relative to all microglial cells in a given mouse are shown. **C.** The microglial soma volume was significantly increased by NIS. **D.** Iron-supplemented HD mice have a significantly increased number of processes relative to vehicle-treated HD mice. **E.** The total process length was significantly decreased in HD mice relative to WT mice but was not changed by iron supplementation. **B-F.** n=10 WT control, 11 WT NIS, 12 HD control, 11 HD NIS. Data are shown as the mean ± SE. *p<0.05, **p<0.01.

### Labile iron accumulates in HD microglia

HD microglia accumulate ferritin-bound iron, which is considered non-pathogenic but is elevated in response to increased labile iron (Simmons Danielle et al., 2007, Picard et al., 1998, Konijn et al., 1999). It is labile iron that has the potential to drive oxidative and inflammatory processes. We used a flow cytometric approach to identify brain microglial cells (Mizutani et al., 2012) and measure cytoplasmic labile iron using the fluorescent dye calcein AM. HD mice had increased labile iron in microglial cells (F_(1,24)_ = 4.67, p = 0.0409), but there was no effect of NIS on labile iron accumulation (**Fig. 5A,B**). To assess whether iron promoted increased generalized neuroinflammation, we quantified expression of the gene encoding kynurenine monooxygenase (*Kmo*) and the chemokine CCL2 (*Ccl2*). The KMO enzyme is downstream of IDO in the KP, and its upregulation favors production of neurotoxic pathway intermediates including 3-hydroxykynurenine and quinolinic acid, which are associated with neuroinflammation (Schwarcz et al., 2012). The chemokine CCL2 (MCP-1), which is expressed by astrocytes and some neurons but not microglia, activates microglial cells and recruits them to sites of inflammation in the central nervous system (Mizutani et al., 2012, Conductier et al., 2010, Hinojosa et al., 2011). Therefore, increased CCL2 is an indicator of a generalized, sustained neuroinflammatory response. *Kmo* transcripts were increased both in HD mice (F_(1,38)_ = 11.29, p = 0.0018) and by NIS (F_(1,38)_ = 5.65, p = 0.0226) in the striatum and cortex (**Fig. 6A,B**). In the striatum, non-supplemented HD mice showed an increase in *Kmo* transcripts compared with WT mice (p = 0.0447) and, in the cortex, iron-supplemented HD mice had increased levels of *Kmo* transcripts compared with non-supplemented HD mice (p = 0.007) and iron-supplemented WT mice (p = 0.0015). No differences in *Ccl2* expression were found in the striatum or cortex (**Fig. 6C,D**)

**Figure 5:**
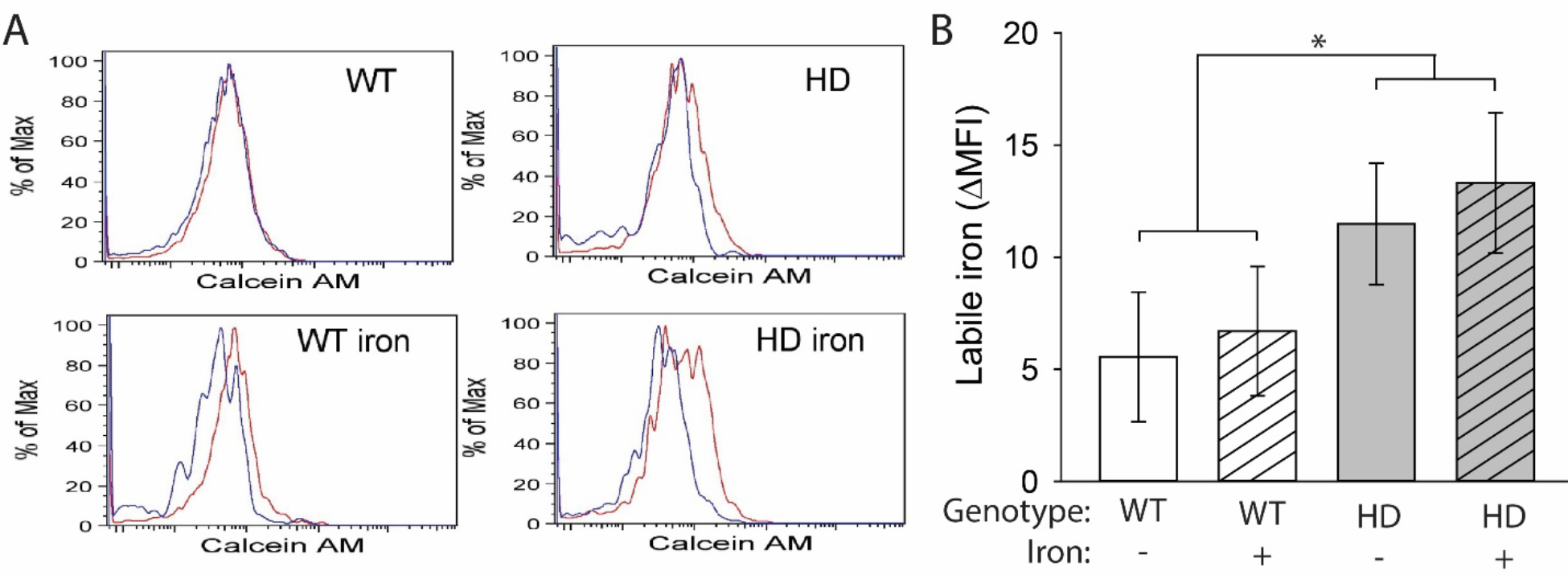
Labile iron accumulates in HD microglia. Microglia, defined as CD11b^+^CD45^+^CX3CR1^+^ cells, were extracted from mouse brains at 14 weeks of age and labeled with calcein AM. **A**. Representative histograms of calcein AM fluorescence in brain microglia incubated with (red) or without (blue) deferiprone. **B**. Labile iron, defined as the difference in mean fluorescence intensity (MFI) between cells incubated with and without deferiprone, was significantly increased in HD microglia. n=6–8. Data are shown as the mean ± SE. *p<0.05.

**Figure 6:**
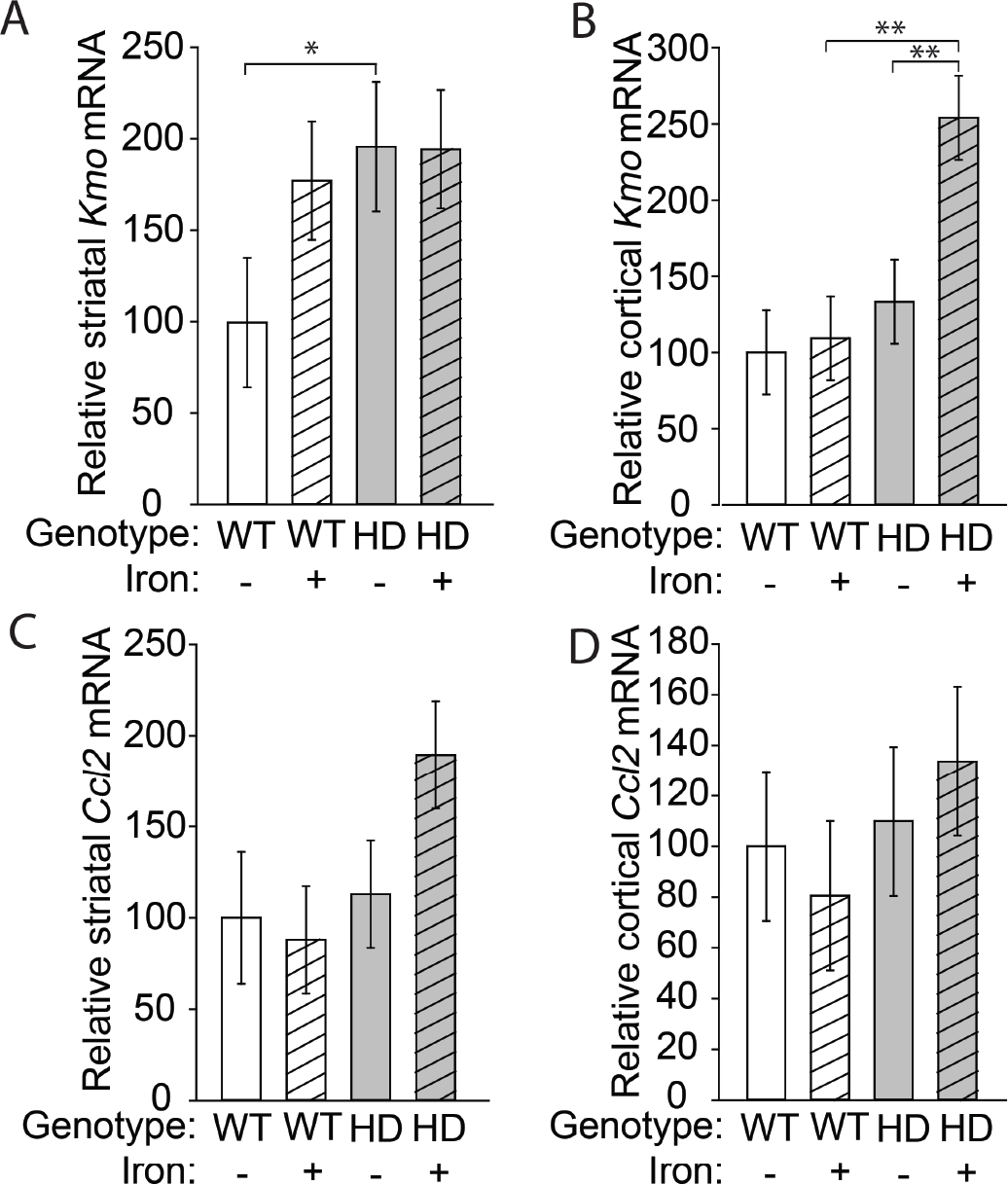
*Kmo* transcripts are increased by HD and NIS. Gene expression was determined at 14 weeks of age. **A**. *Kmo* expression was increased in the mouse HD striatum relative to WT but NIS had not effect. **B.** *Kmo* expression in iron-supplemented HD mice was significantly increased in the cortex. **C-D**. *Ccl2* expression was unchanged in the striatum (**C**) and cortex (**D**). Data are shown as the mean ± SE. n=6.

### Kynurenine metabolism is increased by NIS and HD

We examined KP metabolite profiles to determine if pathway activity downstream of IDO was modified as a result of NIS and / or HD. We found no changes in tryptophan levels among groups in either the cortex or striatum (**Fig. 7A,E**). Kynurenine levels were increased by both neonatal iron (F_(1,80)_ = 5.23, p = 0.0249) and the HD genotype (F_(1,80)_ = 9.95, p = 0.0023) in the cortex and striatum, consistent with IDO activation (**Fig. 7B,F**). We also found a significant effect of both neonatal iron (F_(1,80)_ = 4.23, p = 0.043) and the HD genotype (F_(1,80)_ = 12.7, p = 0.0006) on 3-hydroxykynurenine levels in the cortex and striatum (**Fig. 7C,G**). Although we did not detect an effect of iron supplementation or the HD genotype alone on kynurenic acid levels in either the cortex or striatum, we did find a significant NIS-HD interaction (F_(1,80)_ = 4.06, p = 0.0472) (**Fig. 7D,H**). We also compared metabolite ratios to address the balance of kynurenine metabolism (**Fig. 7I-L**). The ratio of 3-hydroxykynurenine (3-HK) to kynurenic acid is important in determining the relative toxicity of 3-HK, with an increase in the 3-HK/kynurenic acid ratio indicating an increase in the production of neurotoxic kynurenine metabolites (Campesan et al., 2011a). In both the cortex and striatum, we found that 3-HK/kynurenic acid ratios were increased in HD mice (F_(1,40)_ = 7.07, p = 0.0112) but showed only a trend toward being increased by iron supplementation (F_(1,40)_ = 3.09, p = 0.0864) (**Fig. 7K,L**). Interestingly, we found that, in the cortex, iron-supplemented HD mice had an increased 3-HK/kynurenic acid ratio compared with iron-supplemented WT mice (p=0.0031) (**Fig. 7K**). However, with respect to changes in metabolite levels, we did not find differences between iron-supplemented HD mice relative to control HD mice.

**Figure 7:**
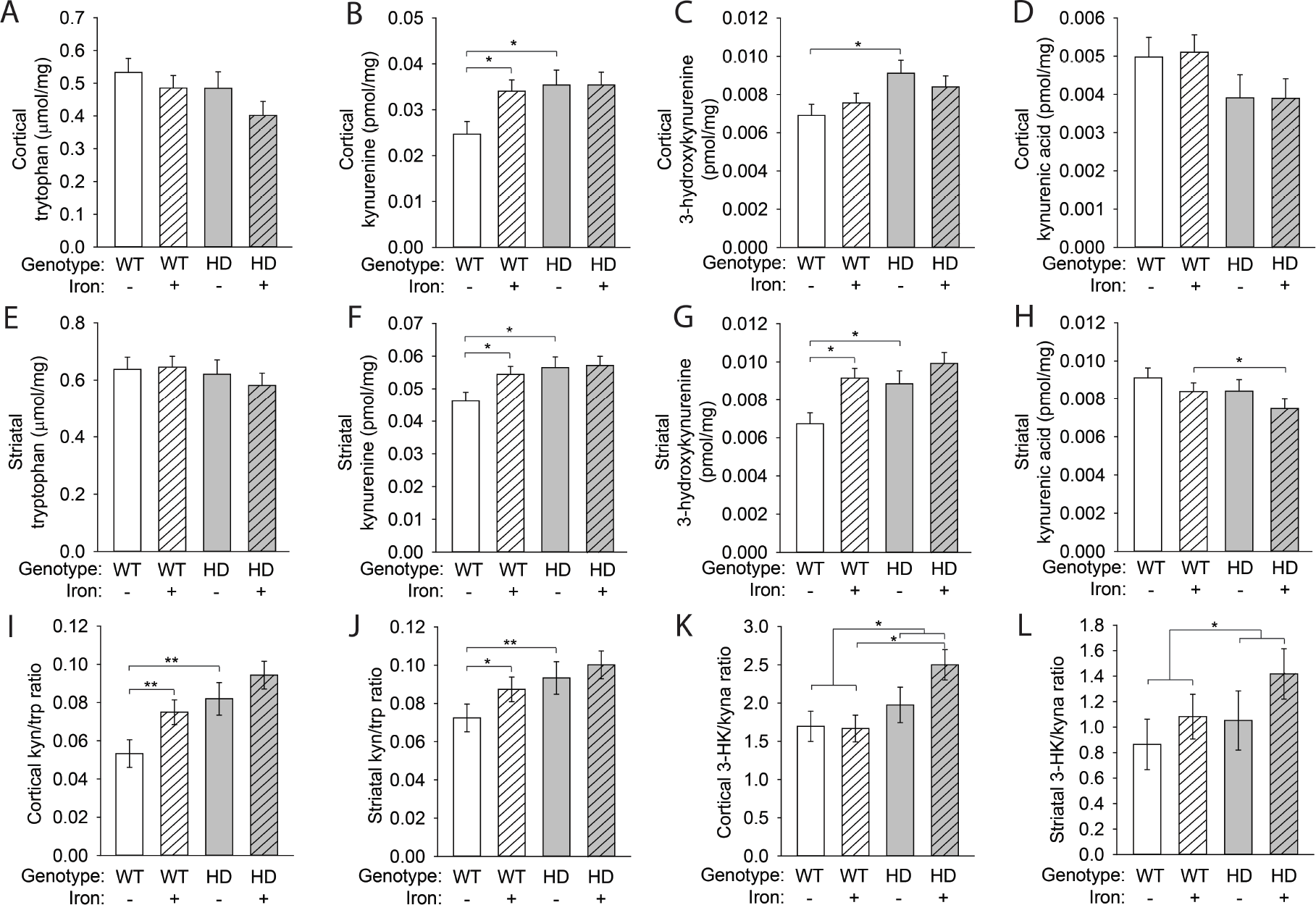
Brain kynurenine metabolism is altered by the HD genotype and NIS. Mouse pups were supplemented with iron from postnatal days 10–17. KP metabolite levels were measured in the cortex (**A-D**) and striatum (**E-H**) at 14 weeks of age and are presented per milligram of soluble brain protein. **A.** Cortical tryptophan levels were unchanged. **B.** Cortical kynurenine was increased in iron-supplemented WT mice and in HD mice relative to control WT mice. **C.** HD mice had elevated 3-HK in the cortex. **D.** Kynurenic acid levels in the cortex were unchanged as a result of iron supplementation or the HD genotype. **E.** Striatal tryptophan levels were unchanged. **F.** Kynurenine was increased in the striatum of iron-supplemented WT mice and in HD mice relative to control WT mice. **G.** Striatal 3-HK was elevated by iron supplementation in WT mice and by the HD genotype **H.** Iron-supplemented HD mice had decreased striatal kynurenic acid compared with iron-supplemented WT mice. **I.** Cortical kynurenine/tryptophan (kyn/trp) ratios were increased in iron-supplemented WT and in HD mice. **J.** The kyn/trp ratio in the striatum was increased by iron supplementation and in HD mice relative to control WT mice. **K.** Iron-supplemented HD mice had increased 3-hydroxykynurenine/kynurenic acid (3-HK/kyna) ratios in the cortex compared with iron-supplemented WT mice. **L.** Striatal 3-HK/kyna ratios were increased by the HD only in the presence of iron supplementation. **A-L.** Data are shown as the mean ± SE. n=10.

## Discussion

Microglial cells are activated early in the course of human HD (Sapp et al., 2001) in association with ferritin accumulation, suggesting a protective mechanism in response to iron accumulation (Simmons Danielle et al., 2007). Studies using iron-selective chelators suggest that iron can be pathogenic in the brain by promoting inflammation. For example, iron chelation prevents neuroinflammation and microglial activation in models of postoperative cognitive impairment (Li et al., 2016, Zhang et al., 2015). However, it has been unclear until now if iron promotes microglial activation and inflammation in HD. The present study demonstrates that labile iron accumulates in HD microglial cells. We show that iron exposure resulting from NIS leads to heightened microglial activation in HD mice, which is associated with increased degeneration. Furthermore, iron directly activates the microglial KP enzyme IDO, providing one mechanism for its HD-promoting effects.

### Neonatal iron supplementation activates microglial cells

To determine a possible role for iron in microglial activation, we used the NIS model. This is relevant to understanding the mechanisms through which iron promotes HD and other disorders because the doses approximate what a human infant may receive and there is no detectable short-term toxicity (Agrawal et al., 2017). We have previously reported that NIS potentiates HD in the R6/2 and YAC128 mouse models (Berggren et al., 2015, Berggren et al., 2016). Consistent with those earlier findings, we now demonstrate that NIS also potentiates disease in the N171-82Q mouse model. Microglial activation results in a graded transition of morphologic changes from ramified cells, which have many processes and small somas, to phagocytic cells, with their large somas and short processes (Torres-Platas et al., 2014). We used blinded qualitative (categorical) and quantitative (continuous) methods in parallel to assess microglial morphology. Consistent with prior reports in the R6/2 mouse and human HD (Simmons Danielle et al., 2007), we demonstrated here that N171-82Q HD mice have morphologic evidence of microglial activation and iron supplementation resulted in additional microglial activation in HD mice. Using a combination of categorical and continuous microglial measurements, we detected a more mild activation in iron-supplemented WT mice relative to NIS in HD mice. The categorical assignment of brain microglia (ramified, primed, amoeboid, reactive) is the sum of several subtle changes including increased soma size, a change in the number of processes, and an overall decrease in process length. Whereas NIS led to increased soma volume, which is one indicator of activation, it did not change the categorical assignment of microglia in WT mice. However, iron-supplemented HD mice displayed increased soma volume (resulting from NIS) as well as decreased process length (due to HD) and had increased microglial activation according to logistic regression analysis. Therefore, together these data suggest that microglial activation in iron-supplemented HD mice results from a synergism between iron and HD that is suggestive of a stronger activation phenotype than is explained by NIS or HD alone.

Cellular iron status is sensed by the central regulators of iron metabolism, iron-regulatory proteins 1 and 2 (IRP1/IRP2) (Klausner et al., 1993). Reports of decreased IRP1/IRP2 combined with increased microglial ferritin are consistent with a response to iron stress in HD mice (Simmons Danielle et al., 2007, Chen et al., 2013). However, it is unclear if these protective responses are completely effective. Iron stress results from labile iron rather than iron accumulation resulting from increased levels of iron-containing proteins. Labile iron can generate hydroxyl radicals via Fenton chemistry, thereby modifying iron-sensitive enzymes (Wang et al., 2016). By isolating brain microglial cells and using flow cytometry, we have demonstrated for the first time that HD microglia have increased labile iron. This finding supports the presence of iron stress in HD microglial cells. We did not, however, find effects of NIS on labile iron. The calcein AM ester used to quantify labile iron is considered mainly a cytoplasmic iron marker. Cytoplasm is a major sources of esterases, which trap most of the calcein after cleavage of the acetoxy-methyl group before it reaches other organelles (Tenopoulou et al., 2007). Therefore, the absence of a detectable increase in labile iron with NIS does not exclude the possibility that there may be a microglial labile iron pool that is not detected by calcein-AM. In this context, we have recently shown, using a different approach, that labile iron accumulates in brain mitochondria from HD mice (Agrawal et al., 2018).

### Labile iron directly activates IDO

The kynurenine pathway of tryptophan degradation is involved in the progression of HD and other brain disorders (Schwarcz et al., 2012). Activation of this pathway occurs early in human HD (Beal et al., 1990, Guidetti et al., 2000). In addition, inhibition of the quinolinic acid branch of this pathway provides protection in HD model systems (Zwilling et al., 2011, Giorgini et al., 2005). IDO is a cytoplasmic enzyme that is expressed in microglia. It catalyzes the first step in this pathway, oxidizing tryptophan to kynurenine, and therefore has an important role in regulating overall pathway activity. We have previously shown that IDO activity is increased in N171-82Q HD mice and that additional activation occurs in the presence of a neuroinflammatory stimulus (Donley et al., 2016). A previous study showed that 3-hydroxyanthranilic acid dioxygenase, a downstream KP enzyme, is activated by iron (Stachowski and Schwarcz, 2012). Consistent with KP modulation, iron activated IDO in cell culture and brain homogenates. Furthermore, *in vivo* activation by NIS in HD mice was decreased by iron chelation *ex vivo*. Importantly, we demonstrated that iron directly activates recombinant human IDO. The labile iron pool has been reported to be as high as 100–300 nM in mammalian cells, but it fluctuates dramatically based on the cell type and environment (Breuer et al., 1996, Konijn et al., 1999, Epsztejn et al., 1997). It is likely that cytosolic labile iron concentrations are lower than these values, but we found that iron activates IDO with an EC_50_ = 1.24 nM. Therefore, our values are well within the physiological range for iron activation to occur in cells.

KP activation is part of the inflammatory response. For example, *Ido* expression is driven by pro-inflammatory cytokines, primarily IFNγ (Yadav et al., 2007). As iron has pro-inflammatory effects, we also determined if there is genetic upregulation of *Ido* with NIS (Kroner et al., 2014, Zhang et al., 2015). Consistent with our prior study, we found increased cortical *Ido1* in non-supplemented N171-82Q HD mice, but there was no effect of NIS (Donley et al., 2016). Furthermore, the role of direct activation of IDO by iron was supported by showing that translational inhibition in cultured microglial cells did not decrease iron-stimulated IDO activity. Therefore, gene upregulation explains some but not all of the increased IDO activity present in HD mice. Together, these data are consistent with a model whereby IDO activity in HD microglia results from basal gene upregulation by inflammatory signaling and direct activation by labile iron. IDO is regulated at the post-translational level; nitric oxide inhibits activity and phosphorylation targets the protein for clearance (Fujigaki et al., 2012, Thomas et al., 2007). However, to our knowledge, this is the first report of a direct activator of IDO. Previous structural analyses have identified allosteric sites on IDO, and, based on our data showing that IDO activity is decreased by iron chelation in brain tissue, it is possible that iron acts at one of these or at another unidentified allosteric site to modulate activity (Lewis-Ballester et al., 2017, Pearson et al., 2010). The mechanism by which IDO oxidizes tryptophan is debated. However, one prevailing hypothesis supported by empirical evidence states that tryptophan oxidation is accomplished through stepwise addition of oxygen molecules followed by a ring-break (Booth et al., 2015, Capece et al., 2012). Therefore, it is also possible that free iron around the active site supports or contributes to the ferryl oxygen and/or the tryptophan-epoxide intermediates during the nucleophilic addition of the second oxygen.

### Neonatal iron supplementation activates kynurenine metabolism

KMO catalyzes the conversion of kynurenine to 3-HK in the first step of the quinolinic branch of the KP. Quinolinic acid and 3-HK can be neurotoxic, and thus KMO activation is thought to be disease promoting. We observed increased *Kmo* transcripts and increases in its catalytic product 3-HK in the cortex of HD mice that received NIS, indicating that enzymatic activity may also be upregulated. The chemokine CCL2 is expressed in astrocytes and peripheral immune cells; it has a role in the activation and recruitment of immune cells, including microglia (Conductier et al., 2010, Hinojosa et al., 2011). In contrast to *Kmo*, *Ccl2* transcript levels were not significantly altered by the HD genotype or NIS, suggesting that there is neither generalized astroglial activation nor monocytic infiltration. HD mice showed changes in brain KP metabolite profiles characterized by elevations in kynurenine and 3-HK. NIS potentiated these increases. Surprisingly, however, this occurred only in WT mice; there was no effect in HD mice. As synthesis and metabolism determine steady-state metabolite levels it is possible that there may be upregulation of other enzymes in this pathway resulting in increased pathway flux. Despite this, our studies show that iron challenge in the form of NIS promotes activation of IDO and an activated microglial morphology consistent with a profound change in microglial activity and function.

Individuals with HD have a pro-inflammatory macrophage phenotype and elevated peripheral TNFα (Di Pardo et al., 2013, Björkqvist et al., 2008). During spinal cord injury, iron loading in macrophages stimulates a switch to a pro-inflammatory phenotype characterized in part by TNFα production (Kroner et al., 2014). In addition, iron accumulation in microglia upregulates matrix metalloproteases that drive neuroinflammation, including MMP-9 (Mairuae et al., 2011). In patients with HD, MMP-9 elevation is correlated with disease burden, and, in mouse models of HD, higher levels of MMP-9 increases the vulnerability of the striatum to injury (J. et al., 2011, Naphade et al., 2017, Connolly et al., 2016). Although our study focused on microglial morphology and the KP, there is thus additional evidence that iron accumulation in microglia may modulate cytokine signaling and other inflammatory pathways. Additionally, IDO activation is implicated in progression of some cancers demonstrating that this work has broad application (Bilir and Sarisozen, 2017, Ferns et al., 2015, Suzuki et al., 2010).

## Materials and Methods

### Materials

All reagents were from Sigma unless otherwise stated.

### Mouse husbandry and breeding

N171-82Q Huntington’s disease mice (Jackson Labs, Bar Harbor, ME; strain B6C3-Tg(HD82Gln)81Gschi/J) were maintained as described (Donley et al., 2016). Sentinel mice were evaluated every 6 months by comprehensive serology panel screening for murine infectious diseases at Charles River Laboratories and were free of all tested diseases. Mice were humanely sacrificed using B-euthanasia solution. This study and all procedures used were approved by the University of Wyoming Institutional Animal Care and Use Committee (UW-IACUC) (protocol number 20150813JF00190-01) in accordance with the National Institutes of Health guidelines.

### Experimental Design

N171-82Q HD mice were used for all experiments. The N171-82Q mouse is a transgenic model of HD that expresses the N-terminal 171 amino acids of human huntingtin protein with 82 poly-glutamine repeats under the control of the prion protein promotor (Schilling et al., 2007). When both males and females were used for experiments, the sex of the mice was included in the statistical analysis. Mouse pups received carbonyl iron supplementation or vehicle control by oral gavage from postnatal day 10–17 as described (Berggren et al., 2015, Berggren et al., 2016). HD and WT littermate mice were systematically assigned to experimental groups at weaning to balance their ages and sex and to minimize effects of litter of origin. Experiments were set up with a 2 × 2 factorial design with genotype and iron treatment as the factors. All researchers were blinded to the genotype and treatment group at weaning and remained blinded until data analysis. All experimental animals are accounted for in the data presented with the exception of three spontaneous mouse deaths. One iron-supplemented WT mouse (at 7 weeks of age) and two control HD mice (at 11 and 13 weeks of age) died during the reported studies.

### Brain indoleamine-2,3-dioxygenase activity

Dissected brain regions were snap frozen on dry ice and stored at −70°C until analysis. Samples were prepared and IDO activity was determined by HPLC-MS/MS as described (Donley et al., 2016). Additionally, a separate aliquot was analyzed the same way but with addition of deferoxamine to the incubation buffer.

### Brain gene expression by real-time quantitative PCR analysis

Total RNA was extracted from the cerebral cortex and striatum using phenol/chloroform and then was purified using a column-based method (RNAeasy, Qiagen) with on-column DNAse 1 digestion. The cDNA was prepared, and *Ido* expression was quantified as described (Donley et al., 2016). *Kmo* and *Ccl2* expression was analyzed using Applied Biosystems Taqman gene expression primer/probe combinations Mm1321343_m1 (*Kmo*) and Mm00441242_m1 (*Ccl2*). Expression was normalized to beta-actin using the Applied Biosystems Taqman gene expression primer/probe combination Mm00607939_s1. Gene expression was determined using 20 ng of cDNA per reaction.

### Purified Indoleamine-2,3-dioxygenase activity

Recombinant human IDO (Accession #P14902) was obtained from R&D Systems (#6030-AO-010) and frozen at −80°C until use. The incubation buffer was prepared by mixing 800 μM tryptophan, 40 mM ascorbic acid, 9000 U/ml catalase, and 40 μM methylene blue in HBSS (pH = 6.5) with 80 mM ascorbic acid in 0.405 M Tris (pH=8.0) in equal parts and pre-warmed to 37°C. One hundred ng/ml purified protein (100 ng/ml) in HBSS was mixed in equal parts with the incubation buffer and incubated at 37°C for 10 minutes. The reaction was stopped by the addition of 20% (v/v) 5 M acetic acid and incubated at 50°C for 10 minutes. Samples were centrifuged at 12,000×g, 4°C for 10 minutes, and the supernatants were filtered through a 0.2-μm filter. Kynurenine was measured in the filtered supernatant as described (Donley et al., 2016).

### Indoleamine-2,3-dioxygenase activity in cultured cells

The EOC20 mouse microglial cell line (ATCC CTRL-2469) was used for experiments. Cells were cultured according to the manufacturer’s instructions and were verified to be mycoplasma free. For IDO measurements, cells were washed twice with prewarmed HBSS and then incubated with IDO incubation buffer as described (Donley et al., 2016) for 4 hours. The kynurenine concentration was quantified in the cell supernatant using the method described above. Cells were lysed, and protein levels quantified using a Bradford assay. IDO activity was defined as picomoles of kynurenine produced per microgram of protein per hour.

### Brain kynurenine metabolite quantification

Dissected brain tissue was snap frozen on dry ice and stored at −80°C until analysis. Brain regions were prepared according to the protocol for IDO activity in brain tissue. Samples were prepared by mixing 450 μg brain protein 1:2 (v/v) with methanol plus 2% acetic acid and then centrifuging the mixture at 12,000×g for 10 minutes at 4°C. Supernatants were filtered through a Phenomenex Phree Phospholipid extraction column and a 0.2-μm filter. The samples were then run on a Waters Acquity UPLC-MS/MS with a methanol/2% acetic acid mobile phase gradient on a 2 × 100–mm Waters BEH C^18^ column. The sample (5 μl) was injected with a total flow rate of 0.3 ml/min and a total run time of 4.5 minutes. Analytes were verified using two M+H parent-daughter transitions. Tryptophan parent m/z = 205.13 and daughter m/z = 118.03 and 146.07; kynurenic acid parent m/z = 190.07 and daughter m/z = 88.96 and 116.03; kynurenine parent m/z = 209.04 and daughter m/z = 94.02 and 146.03; 3-HK parent m/z = 224.98 and daughter m/z = 110.04 and 162.05. Concentrations were quantified using QuanLynx (Waters) software from serial dilutions of standards.

### Mouse neuropathology

Mice designated for stereological analysis were perfused with 4% paraformaldehyde and treated as described (Berggren et al., 2015, Berggren et al., 2016). Briefly, fixed brains were sectioned at 40 μm on a freezing microtome, and serial sections were collected in 12-well plates so that one well contained every 12^th^ brain section. Sections from one well were mounted and thionin stained. The Cavalieri method and nucleator methods in StereoInvestigator (MicroBrightField) software were used to determine striatal volume and neuronal cell body volume, respectively.

### Microglial immunohistochemistry

Microglial morphology was characterized in the striatum at the level of the anterior commissure. Striatal sections were mounted, and microglia were immunolabeled with anti-Iba1 (Abcam ab178846). Brain sections were permeabilized for 15 minutes in methanol + 3% hydrogen peroxide and then blocked with 1× TBE, 0.05% Tween-20, and 2% normal serum. Sections were incubated for 3 days in primary antibody (1:1000) with appropriate controls in blocking buffer at 4°C. They were then washed and incubated for 24 hours with biotinylated secondary antibody prior to washing, labeling with streptavidin-HRP conjugate (Vector Labs), and detection using 3,3’-diaminobenzidine per the manufacturer’s instructions. Reactions were quenched with water and then washed twice in 1× TBE prior to coverslipping with fluoromount G (Southern Biotech).

### Microglial morphology characterization

Microglial cells were defined as Iba1-positive cells in brain sections. Two 50-μm image stacks were collected from the dorsal and medial portion of the striatum at the level of the anterior commissure. From these images, two independent approaches were used to quantify the microglial morphology. First, cells were classified as ramified, primed, reactive, or amoeboid as described (Torres-Platas et al., 2014). Second, quantitative morphology parameters were determined using Neurolucida software (MicroBrightField). Image stacks were uploaded to Neurolucida, and each cell was traced throughout the stack. All cells that were fully in the image stack were evaluated. Ten microglial cells or more were traced per brain. The tracings were then uploaded to Neurolucida Explorer, where they were analyzed by branched structure analysis.

### Microglial labile iron analysis

We used flow cytometry to identify microglia purified from forebrains in combination with calcein AM detection of labile iron. In brief, mice were perfused for two minutes with cold, heparinized saline. Forebrains were dissected out and placed in cold HBSS and then diced, digested, and homogenized to isolate brain microglia as described (Grabert et al., 2016, Nikodemova and Watters, 2012). Cells from each animal were plated at 1 × 10^6^ cells/ml in four separate wells. Cells were incubated with primary antibody conjugates for 30 minutes. Microglial cells were labeled using CD11b, CD45, and CX3CR1 (Biolegend) and analyzed by flow cytometry. Microglial markers including anti-Iba1 and CD68 were not used at the risk of losing iron from the cell due to the permeabilization required for intracellular staining. Previously, calcein AM was used to characterize the labile iron pool in cells by comparing calcein AM fluorescence with and without an iron chelator added to cells (Ali et al., 2003, Prus and Fibach, 2007). Two wells from each brain were incubated with 10 μM deferiprone, a membrane permeable iron chelator, in HBSS for 30 minutes at room temperature, and two wells received buffer at the same time. Fixable Near-IR Live/Dead Stain was used to exclude dead cells. (ThermoFisher Scientific) To detect iron, the cells were also incubated with 20 μM calcein AM for 15 minutes. Cells were washed thoroughly, and flow cytometry data were collected using a Guava easyCyte HT flow cytometer (Millipore). Data were analyzed using FlowJo v9.3 software. The cell gating and compensation strategy was determined from no-stain, single-color, and fluorescence-minus-one controls for each fluorescence channel. The mean fluorescence intensity was determined for each sample. Labile iron in microglia was defined as the difference in the mean fluorescence intensity for cells from the same mouse that were labeled with calcein AM with and without deferiprone treatment, similar to that described previously (Ali et al., 2003, Prus and Fibach, 2007).

### Statistical analyses

Data were analyzed using SAS software version 9.2 (Cary, NC). Generalized linear modeling was used for single-timepoint analyses, and mixed modeling was used for repeated measures and matched data. Assumptions of normality and equal variance were verified for each experiment. All main effects and interactions were investigated in the initial statistical model, and then we performed pre-planned pairwise comparisons. Non-normally distributed data were log transformed for analysis and are presented as the mean ± 95% CIs, as distributions were significantly right-skewed. Logistic regression was used to analyze the categorical data on microglial morphology and to determine interaction effects between treatment and genotype. Logistic regression yielded an inference on the odds of microglial activation based on an activated microglial morphology. This statistical method is powerful against non-linear effects and for data, such as microglial morphology, where the interval between variables is not constant. Differences were considered significant with p<0.05. Asterisks in figures represent statistical significance resulting from individual comparisons.

## Acknowledgements

The authors would like to thank Dr. Kenneth Gerow, University of Wyoming, for consulting on statistical analysis of microglial morphologic data. Funding for this study was provided by NIH NINDS R01 NS079450. The Wyoming INBRE program, supported by the Institutional Development Award from NIH NIGMS 2P20GM103432, provided a graduate fellowship to DWD and an undergraduate research fellowship to MR. The funding agencies had no input on study design, data analysis, or decision to publish. The authors report no conflicts of interest.

## References

Agrawal, S., Berggren, K. L., Marks, E. & Fox, J. H. 2017. Impact of high iron intake on cognition and neurodegeneration in humans and in animal models: a systematic review. Nutrition Reviews, 75, 456–470.

Agrawal, S., Fox, J., Thyagarajan, B. & Fox, J. 2018. Brain mitochondrial iron accumulates in Huntington’s disease, mediates mitochondrial dysfunction, and can be removed pharmacologically. Free Radical Biology and Medicine.

Ali, A., Zhang, Q., Dai, J. & Huang, X. 2003. Calcein as a fluorescent iron chemosensor for the determination of low molecular weight iron in biological fluids. Biometals, 16, 285–293.

Andersen, H. H., Johnsen, K. B. & Moos, T. 2014. Iron deposits in the chronically inflamed central nervous system and contributes to neurodegeneration. Cellular and Molecular Life Sciences, 71, 1607–1622.

Ball, H. J., Sanchez-Perez, A., Weiser, S., Austin, C. J. D., Astelbauer, F., Miu, J., Mcquillan, J. A., Stocker, R., Jermiin, L. S. & Hunt, N. H. 2007. Characterization of an indoleamine 2,3-dioxygenase-like protein found in humans and mice. Gene, 396, 203–213.

Bartzokis, G., Cummings, J., Perlman, S., Hance, D. B. & Mintz, J. 1999. Increased basal ganglia iron levels in huntington disease. Archives of Neurology, 56, 569–574.

Beal, M. F., Matson, W. R., Swartz, K. J., Gamache, P. H. & Bird, E. D. 1990. Kynurenine pathway measurements in Huntington’s disease striatum: evidence for reduced formation of kynurenic acid. J Neurochem, 55, 1327–39.

Berggren, K. L., Chen, J., Fox, J., Miller, J., Dodds, L., Dugas, B., Vargas, L., Lothian, A., Mcallum, E., Volitakis, I., Roberts, B., Bush, A. I. & Fox, J. H. 2015. Neonatal iron supplementation potentiates oxidative stress, energetic dysfunction and neurodegeneration in the R6/2 mouse model of Huntington’s disease. Redox Biology, 4, 363–374.

Berggren, K. L., Lu, Z., Fox, J. A., Dudenhoeffer, M., Agrawal, S. & Fox, J. H. 2016. Neonatal Iron Supplementation Induces Striatal Atrophy in Female YAC128 Huntington’s Disease Mice. J Huntingtons Dis, 5, 53–63.

Bilir, C. & Sarisozen, C. 2017. Indoleamine 2,3-dioxygenase (IDO): Only an enzyme or a checkpoint controller? Journal of Oncological Sciences, 3, 52–56.

Björkqvist, M., Wild, E. J., Thiele, J., Silvestroni, A., Andre, R., Lahiri, N., Raibon, E., Lee, R. V., Benn, C. L., Soulet, D., Magnusson, A., Woodman, B., Landles, C., Pouladi, M. A., Hayden, M. R., Khalili-Shirazi, A., Lowdell, M. W., Brundin, P., Bates, G. P., Leavitt, B. R., Möller, T. & Tabrizi, S. J. 2008. A novel pathogenic pathway of immune activation detectable before clinical onset in Huntington’s disease. The Journal of Experimental Medicine, 205, 1869–1877.

Booth, E. S., Basran, J., Lee, M., Handa, S. & Raven, E. L. 2015. Substrate Oxidation by Indoleamine 2,3-Dioxygenase: Evidence for a Common Reaction Mechanism. Journal of Biological Chemistry.

Breuer, W., Epsztejn, S. & Cabantchik, Z. I. 1996. Dynamics of the cytosolic chelatable iron pool of K562 cells. FEBS Lett, 382, 304–8.

Campesan, S., Green, EDWARD W., Breda, C., Sathyasaikumar, KORRAPATI V., Muchowski, PAUL J., Schwarcz, R., Kyriacou, CHARALAMBOS P. & Giorgini, F. 2011a. The Kynurenine Pathway Modulates Neurodegeneration in a Drosophila Model of Huntington’s Disease. Current Biology, 21, 961–966.

Campesan, S., Green, E. W., Breda, C., Sathyasaikumar, K. V., Muchowski, P. J., Schwarcz, R., Kyriacou, C. P. & Giorgini, F. 2011b. The kynurenine pathway modulates neurodegeneration in a Drosophila model of Huntington’s disease. Curr Biol, 21, 961–6.

Capece, L., Lewis-Ballester, A., Yeh, S.-R., Estrin, D. A. & Marti, M. A. 2012. Complete Reaction Mechanism of Indoleamine 2,3-Dioxygenase as Revealed by QM/MM Simulations. The Journal of Physical Chemistry B, 116, 1401–1413.

Castelnau, P. A., Garrett, R. S., Palinski, W., Witztum, J. L., Campbell, I. L. & Powell, H. C. 1998. Abnormal iron deposition associated with lipid peroxidation in transgenic mice expressing interleukin-6 in the brain. J Neuropathol Exp Neurol, 57, 268–82.

Chen, J., Marks, E., Lai, B., Zhang, Z., Duce, J. A., Lam, L. Q., Volitakis, I., Bush, A. I., Hersch, S. & Fox, J. H. 2013. Iron Accumulates in Huntington’s Disease Neurons: Protection by Deferoxamine. PLOS ONE, 8, e77023.

Conductier, G., Blondeau, N., Guyon, A., Nahon, J. L. & Rovere, C. 2010. The role of monocyte chemoattractant protein MCP1/CCL2 in neuroinflammatory diseases. J Neuroimmunol, 224, 93–100.

Connolly, C., Magnusson-Lind, A., Lu, G., Wagner, P. K., Southwell, A. L., Hayden, M. R., Björkqvist, M. & Leavitt, B. R. 2016. Enhanced immune response to MMP3 stimulation in microglia expressing mutant huntingtin. Neuroscience, 325, 74–88.

Corona, A. W., Norden, D. M., Skendelas, J. P., Huang, Y., O’connor, J. C., Lawson, M., Dantzer, R., Kelley, K. W. & Godbout, J. P. 2013. Indoleamine 2,3-dioxygenase inhibition attenuates lipopolysaccharide induced persistent microglial activation and depressive-like complications in fractalkine receptor (CX3CR1)-deficient mice. Brain, Behavior, and Immunity, 31, 134–142.

Davis, B. M., Salinas-Navarro, M., Cordeiro, M. F., Moons, L. & De groef, L. 2017. Characterizing microglia activation: a spatial statistics approach to maximize information extraction. Scientific Reports, 7, 1576.

Dexter, D. T., Carayon, A., Javoy-Agid, F., Agid, Y., Wells, F. R., Daniel, S. E., Lees, A. J., Jenner, P. & Marsden, C. D. 1991. Alterations in the levels of iron, ferritin and other trace metals in Parkinson’s disease and other neurodegenerative diseases affecting the basal ganglia. Brain, 114 (Pt 4), 1953–75.

Di pardo, A., Alberti, S., Maglione, V., Amico, E., Cortes, E. P., Elifani, F., Battaglia, G., Busceti, C. L., Nicoletti, F., Vonsattel, J. P. G. & Squitieri, F. 2013. Changes of peripheral TGF-β1 depend on monocytes-derived macrophages in Huntington disease. Molecular Brain, 6, 55–55.

Dietrich, P., Johnson, I. M., Alli, S. & Dragatsis, I. 2017. Elimination of huntingtin in the adult mouse leads to progressive behavioral deficits, bilateral thalamic calcification, and altered brain iron homeostasis. PLOS Genetics, 13, e1006846.

Donley, D. W., Olson, A. R., Raisbeck, M. F., Fox, J. H. & Gigley, J. P. 2016. Huntingtons Disease Mice Infected with Toxoplasma gondii Demonstrate Early Kynurenine Pathway Activation, Altered CD8+ T-Cell Responses, and Premature Mortality. PLOS ONE, 11, e0162404.

Epsztejn, S., Kakhlon, O., Glickstein, H., Breuer, W. & Cabantchik, Z. I. 1997. Fluorescence Analysis of the Labile Iron Pool of Mammalian Cells. Analytical Biochemistry, 248, 31–40.

Fernández-arjona, M. D. M., Grondona, J. M., Granados-durán, P., Fernández-llebrez, P. & López-ávalos, M. D. 2017. Microglia Morphological Categorization in a Rat Model of Neuroinflammation by Hierarchical Cluster and Principal Components Analysis. Frontiers in Cellular Neuroscience, 11.

Ferns, D. M., Kema, I. P., Buist, M. R., Nijman, H. W., Kenter, G. G. & Jordanova, E. S. 2015. Indoleamine-2,3-dioxygenase (IDO) metabolic activity is detrimental for cervical cancer patient survival. OncoImmunology, 4, e981457.

Fujigaki, H., Seishima, M. & Saito, K. 2012. Posttranslational modification of indoleamine 2,3-dioxygenase. Analytical and Bioanalytical Chemistry, 403, 1777–1782.

Garcia-Miralles, M., Hong, X., Tan, L. J., Caron, N. S., Huang, Y., To, X. V., Lin, R. Y., Franciosi, S., Papapetropoulos, S., Hayardeny, L., Hayden, M. R., Chuang, K.-H. & Pouladi, M. A. 2016. Laquinimod rescues striatal, cortical and white matter pathology and results in modest behavioural improvements in the YAC128 model of Huntington disease. Scientific Reports, 6, 31652.

Giorgini, F., Guidetti, P., Nguyen, Q., Bennett, S. C. & Muchowski, P. J. 2005. A genomic screen in yeast implicates kynurenine 3-monooxygenase as a therapeutic target for Huntington’s disease. Nature genetics, 37, 526–531.

Grabert, K., Michoel, T., Karavolos, M. H., Clohisey, S., Baillie, J. K., Stevens, M. P., Freeman, T. C., Summers, K. M. & Mccoll, B. W. 2016. Microglial brain region–dependent diversity and selective regional sensitivities to aging. Nature Neuroscience, 19, 504.

Guidetti, P., Hemachandra Reddy, P., Tagle, D. A. & Schwarcz, R. 2000. Early kynurenergic impairment in Huntington’s Disease and in a transgenic animal model. Neuroscience Letters, 283, 233–235.

Guidetti, P., Luthi-Carter, R. E., Augood, S. J. & Schwarcz, R. 2004. Neostriatal and cortical quinolinate levels are increased in early grade Huntington’s disease. Neurobiology of Disease, 17, 455–461.

H.P., J. A., Maurik, V. H., C., O. D. K. I., T., M. R., Anna-Aster, D. R., H., S. M., L., S. D., Corien, G., Nieske, B., Willem, K., H.W.G.M., B., F.A., D. D. W., M.C., V. R. W., P., B. G., M., H. E. & A., R. E. 2017. Frequency of nuclear mutant huntingtin inclusion formation in neurons and glia is cell-type-specific. Glia, 65, 50–61.

HDCRG 1993. A novel gene containing a trinucleotide repeat that is expanded and unstable on Huntington’s disease chromosomes. The Huntington’s Disease Collaborative Research Group (HDCRG). Cell, 72, 971–83.

Hinojosa, A. E., Garcia-Bueno, B., Leza, J. C. & Madrigal, J. L. 2011. CCL2/MCP-1 modulation of microglial activation and proliferation. J Neuroinflammation, 8, 77.

J., D. V., J., D. V., G., M., A., C., M., P., J., V. & C., P. 2011. Role of matrix metalloproteinase-9 (MMP-9) in striatal blood–brain barrier disruption in a 3-nitropropionic acid model of Huntington’s disease. Neuropathology and Applied Neurobiology, 37, 525–537.

Jurgens, C. K., Jasinschi, R., Ekin, A., Witjes-Ane, M. N., Middelkoop, H., Van der grond, J. & Roos, R. A. 2010. MRI T2 Hypointensities in basal ganglia of premanifest Huntington’s disease. PLoS Curr, 2.

Kierdorf, K. & Prinz, M. 2013. Factors regulating microglia activation. Frontiers in Cellular Neuroscience, 7.

Klausner, R. D., Rouault, T. A. & Harford, J. B. 1993. Regulating the fate of mRNA: The control of cellular iron metabolism. Cell, 72, 19–28.

Konijn, A. M., Glickstein, H., Vaisman, B., Meyron-Holtz, E. G., Slotki, I. N. & Cabantchik, Z. I. 1999. The Cellular Labile Iron Pool and Intracellular Ferritin in K562 Cells. Blood, 94, 2128–2134.

Kraft, A. D., Kaltenbach, L. S., Lo, D. C. & Harry, G. J. 2012. Activated microglia proliferate at neurites of mutant huntingtin-expressing neurons. Neurobiology of aging, 33, 621.e17–33.

Kroner, A., Greenhalgh, ANDREW D., Zarruk, JUAN G., Passos dos santos, R., Gaestel, M. & David, S. 2014. TNF and Increased Intracellular Iron Alter Macrophage Polarization to a Detrimental M1 Phenotype in the Injured Spinal Cord. Neuron, 83, 1098–1116.

La spada, A. R., Weydt, P. & Pineda, V. V. 2011. Frontiers in Neuroscience Huntington’s Disease Pathogenesis: Mechanisms and Pathways. In: Lo, D. C. & Hughes, R. E. (eds.) Neurobiology of Huntington’s Disease: Applications to Drug Discovery. Boca Raton (FL): CRC Press/Taylor & Francis Llc.

Lawson, L. J., Perry, V. H., Dri, P. & Gordon, S. 1990. Heterogeneity in the distribution and morphology of microglia in the normal adult mouse brain. Neuroscience, 39, 151–70.

Lee, A., Kanuri, N., Zhang, Y., Sayuk, G. S., Li, E. & Ciorba, M. A. 2014. IDO1 and IDO2 Non-Synonymous Gene Variants: Correlation with Crohn’s Disease Risk and Clinical Phenotype. PLOS ONE, 9, e115848.

Lewis-Ballester, A., Pham, K. N., Batabyal, D., Karkashon, S., Bonanno, J. B., Poulos, T. L. & Yeh, S.-R. 2017. Structural insights into substrate and inhibitor binding sites in human indoleamine 2,3-dioxygenase 1. Nature Communications, 8, 1693.

Li, Y., Pan, K., Chen, L., Ning, J.-L., Li, X., Yang, T., Terrando, N., Gu, J. & Tao, G. 2016. Deferoxamine regulates neuroinflammation and iron homeostasis in a mouse model of postoperative cognitive dysfunction. Journal of Neuroinflammation, 13, 268.

Lopes, K. O., Sparks, D. L. & Streit, W. J. 2008. Microglial dystrophy in the aged and Alzheimer’s disease brain is associated with ferritin immunoreactivity. Glia, 56, 1048–60.

Lu, Z., Marks, E., Chen, J., Moline, J., Barrows, L., Raisbeck, M., Volitakis, I., Cherny, R. A., Chopra, V., Bush, A. I., Hersch, S. & Fox, J. H. 2014. Altered selenium status in Huntington’s disease: neuroprotection by selenite in the N17182Q mouse model. Neurobiol Dis, 71, 34–42.

Lumsden, A. L., Henshall, T. L., Dayan, S., Lardelli, M. T. & Richards, R. I. 2007. Huntingtin-deficient zebrafish exhibit defects in iron utilization and development. Human Molecular Genetics, 16, 1905–1920.

Mairuae, N., Connor, J. R. & Cheepsunthorn, P. 2011. Increased cellular iron levels affect matrix metalloproteinase expression and phagocytosis in activated microglia. Neuroscience Letters, 500, 36–40.

Masuda, N., Peng, Q., Li, Q., Jiang, M., Liang, Y., Wang, X., Zhao, M., Wang, W., Ross, C. A. & Duan, W. 2008. Tiagabine is neuroprotective in the N171-82Q and R6/2 mouse models of Huntington’s disease. Neurobiology of disease, 30, 293–302.

Mccarthy, R. C., Sosa, J. C., Gardeck, A. M., Baez, A. S., Lee, C.-H. & Wessling-Resnick, M. 2018. Inflammation-induced Iron Transport and Metabolism by Brain Microglia. Journal of Biological Chemistry.

Mehlhase, J., Gieche, J., Widmer, R. & Grune, T. 2006. Ferritin levels in microglia depend upon activation: Modulation by reactive oxygen species. Biochimica et Biophysica Acta (BBA) - Molecular Cell Research, 1763, 854–859.

Metz, R., Duhadaway, J. B., Kamasani, U., Laury-Kleintop, L., Muller, A. J. & Prendergast, G. C. 2007. Novel Tryptophan Catabolic Enzyme IDO2 Is the Preferred Biochemical Target of the Antitumor Indoleamine 2,3-Dioxygenase Inhibitory Compound <span class=“sc”>d</span>-1-Methyl-Tryptophan. Cancer Research, 67, 7082–7087.

Micronutrients, I. O. M. P. O. 2001. Dietary Reference Intakes for Vitamin A, Vitamin K, Arsenic, Boron, Chromium, Copper, Iodine, Iron, Manganese, Molybdenum, Nickel, Silicon, Vanadium, and Zinc. [Online]. National Academies Press (US). Available: https://www.ncbi.nlm.nih.gov/books/NBK222310/ [Accessed 2018].

Mizutani, M., Pino, P. A., Saederup, N., Charo, I. F., Ransohoff, R. M. & Cardona, A. E. 2012. The fractalkine receptor but not CCR2 is present on microglia from embryonic development throughout adulthood(). Journal of immunology (Baltimore, Md.: 1950), 188, 29–36.

Naphade, S., Embusch, A., Madushani, K. L., Ring, K. L. & Ellerby, L. M. 2017. Altered Expression of Matrix Metalloproteinases and Their Endogenous Inhibitors in a Human Isogenic Stem Cell Model of Huntington’s Disease. Frontiers in Neuroscience, 11, 736.

Nikodemova, M. & Watters, J. J. 2012. Efficient isolation of live microglia with preserved phenotypes from adult mouse brain. Journal of Neuroinflammation, 9, 147.

Pearson, J. T., Siu, S., Meininger, D. P., Wienkers, L. C. & Rock, D. A. 2010. In Vitro Modulation of Cytochrome P450 Reductase Supported Indoleamine 2,3-Dioxygenase Activity by Allosteric Effectors Cytochrome b5 and Methylene Blue. Biochemistry, 49, 2647–2656.

Picard, V., Epsztejn, S., Santambrogio, P., Cabantchik, Z. I. & Beaumont, C. 1998. Role of Ferritin in the Control of the Labile Iron Pool in Murine Erythroleukemia Cells. Journal of Biological Chemistry, 273, 15382–15386.

Prus, E. & Fibach, E. 2007. Flow cytometry measurement of the labile iron pool in human hematopoietic cells. Cytometry Part A, 73A, 22–27.

Rathnasamy, G., Ling, E. A. & Kaur, C. 2011. Iron and iron regulatory proteins in amoeboid microglial cells are linked to oligodendrocyte death in hypoxic neonatal rat periventricular white matter through production of proinflammatory cytokines and reactive oxygen/nitrogen species. J Neurosci, 31, 17982–95.

Rosas, H. D., Chen, Y. I., Doros, G., Salat, D. H., Chen, N. K., Kwong, K. K., Bush, A., Fox, J. & Hersch, S. M. 2012. Alterations in brain transition metals in Huntington disease: an evolving and intricate story. Arch Neurol, 69, 887–93.

Sapp, E., Kegel, K. B., Aronin, N., Hashikawa, T., Uchiyama, Y., Tohyama, K., Bhide, P. G., Vonsattel, J. P. & Difiglia, M. 2001. Early and progressive accumulation of reactive microglia in the Huntington disease brain. J Neuropathol Exp Neurol, 60, 161–72.

Schilling, G., Klevytska, A., Tebbenkamp, A. T., Juenemann, K., Cooper, J., Gonzales, V., Slunt, H., Poirer, M., Ross, C. A. & Borchelt, D. R. 2007. Characterization of huntingtin pathologic fragments in human Huntington disease, transgenic mice, and cell models. J Neuropathol Exp Neurol, 66, 313–20.

Schwarcz, R., Bruno, J. P., Muchowski, P. J. & Wu, H.-Q. 2012. Kynurenines in the mammalian brain: when physiology meets pathology. Nature Reviews Neuroscience, 13, 465.

Simmons danielle, A., Casale, M., Alcon, B., Pham, N., Narayan, N. & Lynch, G. 2007. Ferritin accumulation in dystrophic microglia is an early event in the development of Huntington’s disease. Glia, 55, 1074–1084.

Stachowski, E. K. & Schwarcz, R. 2012. Regulation of quinolinic acid neosynthesis in mouse, rat and human brain by iron and iron chelators in vitro. J Neural Transm (Vienna), 119, 123–31.

Suzuki, Y., Suda, T., Furuhashi, K., Suzuki, M., Fujie, M., Hahimoto, D., Nakamura, Y., Inui, N., Nakamura, H. & Chida, K. 2010. Increased serum kynurenine/tryptophan ratio correlates with disease progression in lung cancer. Lung Cancer, 67, 361–365.

Tai, Y. F., Pavese, N., Gerhard, A., Tabrizi, S. J., Barker, R. A., Brooks, D. J. & Piccini, P. 2007. Microglial activation in presymptomatic Huntington’s disease gene carriers. Brain, 130, 1759–66.

Tenopoulou, M., Kurz, T., Doulias, P.-T., Galaris, D. & Brunk, ULF T. 2007. Does the calcein-AM method assay the total cellular ‘labile iron pool’ or only a fraction of it? Biochemical Journal, 403, 261–266.

Thomas, S. R., Terentis, A. C., Cai, H., Takikawa, O., Levina, A., Lay, P. A., Freewan, M. & Stocker, R. 2007. Post-translational Regulation of Human Indoleamine 2,3-Dioxygenase Activity by Nitric Oxide. Journal of Biological Chemistry, 282, 23778–23787.

Torres-Platas, S. G., Comeau, S., Rachalski, A., Bo, G. D., Cruceanu, C., Turecki, G., Giros, B. & Mechawar, N. 2014. Morphometric characterization of microglial phenotypes in human cerebral cortex. Journal of Neuroinflammation, 11, 12–12.

Van Bergen, J. M. G., Hua, J., Unschuld, P. G., Lim, I. A. L., Jones, C. K., Margolis, R. L., Ross, C. A., Van zijl, P. C. M. & Li, X. 2016. Quantitative susceptibility mapping suggests altered brain iron in premanifest Huntington’s disease. AJNR. American journal of neuroradiology, 37, 789–796.

Vonsattel, J. P. G. & Difiglia, M. 1998. Huntington disease. Journal of Neuropathology and Experimental Neurology, 57, 369–384.

Vymazal, J., Klempíř, J., Jech, R., Židovská, J., Syka, M., Růžička, E. & Roth, J. 2007. MR relaxometry in Huntington’s disease: Correlation between imaging, genetic and clinical parameters. Journal of the Neurological Sciences, 263, 20–25.

Wang, B., Lu, J., Dubey, K. D., Dong, G., Lai, W. & Shaik, S. 2016. How do Enzymes Utilize Reactive OH Radicals? Lessons from Nonheme HppE and Fenton Systems. Journal of the American Chemical Society, 138, 8489–8496.

Wexler, N. S. 2004. Venezuelan kindreds reveal that genetic and environmental factors modulate Huntington’s disease age of onset. Proceedings of the National Academy of Sciences of the United States of America, 101, 3498–3503.

Yadav, M. C., Burudi, E. M. E., Alirezaei, M., Flynn, C. C., Watry, D. D., Lanigan, C. M. & Fox, H. S. 2007. IFN-γ induced IDO and WRS expression in microglia is differentially regulated by IL-4. Glia, 55, 1385–1396.

Yoshida, T., Tanaka, M., Sotomatsu, A. & Hirai, S. 1995. Activated microglia cause superoxide-mediated release of iron from ferritin. Neurosci Lett, 190, 21–4.

Zhang, W., Yan, Z.-F., Gao, J.-H., Sun, L., Huang, X.-Y., Liu, Z., Yu, S.-Y., Cao, C.-J., Zuo, L.-J., Chen, Z.-J., Hu, Y., Wang, F., Hong, J.-S. & Wang, X.-M. 2014. Role and Mechanism of Microglial Activation in Iron-Induced Selective and Progressive Dopaminergic Neurodegeneration. Molecular neurobiology, 49, 1153–1165.

Zhang, X.-Y., Cao, J.-B., Zhang, L.-M., Li, Y.-F. & Mi, W.-D. 2015. Deferoxamine attenuates lipopolysaccharide-induced neuroinflammation and memory impairment in mice. Journal of Neuroinflammation, 12, 20.

Zwilling, D., Huang, S.-Y., Sathyasaikumar, KORRAPATI V., Notarangelo, FRANCESCA M., Guidetti, P., Wu, H.-Q., Lee, J., Truong, J., Andrews-Zwilling, Y., Hsieh, ERIC W., Louie, JAMIE Y., Wu, T., Scearce-Levie, K., Patrick, C., Adame, A., Giorgini, F., Moussaoui, S., Laue, G., Rassoulpour, A., Flik, G., Huang, Y., Muchowski, JOSEPH M., Masliah, E., Schwarcz, R. & Muchowski, PAUL J. 2011. Kynurenine 3-Monooxygenase Inhibition in Blood Ameliorates Neurodegeneration. Cell, 145, 863–874.

